# Activity-dependent lipid droplet biogenesis and turnover regulate synaptic integrity

**DOI:** 10.64898/2026.06.25.734633

**Authors:** Eleni Katafygiotou, Ada Squires, Anne Liang, Holden Hadfield, João A Paulo, Walid Idi, Steven P Gygi, Joongkyu Park, Jeeyun Chung

## Abstract

Lipid droplets (LDs) are conserved organelles that buffer lipid storage and stress, yet their dynamics and functions in neurons remain largely unknown. Here, we report activity-dependent dynamics of neuronal LDs, visualized by a novel, genetically encoded LD reporter (termed LipiDew), in both cultured neurons and mouse motor cortex. Using LipiDew, we found that various paradigms of neuronal activation induced predominant and transient formation of LDs in neurites. Disruption of autophagic LD degradation (lipophagy) resulted in abnormal lipid accumulation in dendritic spines and shafts, promoted recruitment of synaptic scaffolding proteins to LDs, and altered intracellular calcium kinetics in neurons. In addition, mice with neuron-specific genetic impairment of lipophagy showed motor function defects. Together, these findings identify activity-dependent LD formation and lipophagic clearance in neuronal compartments as a crucial regulatory mechanism of synaptic integrity and neuronal function.

## Main Text

Brains require a robust energy supply and membrane plasticity to function. Thus, lipid homeostasis in brain cells is pivotal for governing healthy neural structure and function (*1*, *2*). Subcellular organelles called lipid droplets (LDs) play integral roles in lipid metabolism by storing lipids and buffering fluctuations in cellular fatty acids to prevent lipotoxicity and to provide lipid sources as needed (*3–5*). While LDs in various cell types have been extensively studied (*3*), the dynamics and functions of neuronal LDs remain poorly understood, despite evidence of lipid-protein aggregates in intra- and extraneuronal environments in many neurodegenerative diseases (*1*, *6–10*).

Intriguingly, genes that regulate LD biogenesis and mobilization—including cytosolic lipolysis and lysosomal LD degradation through lipophagy—are associated in humans with hereditary spastic paraplegia (HSP), a neurodegenerative disorder characterized by progressive lower-limb spasticity and weakness (*7*, *11–14*). The lipophagy receptor spartin, encoded by *SPG20*, has been implicated in LD turnover, and loss-of-function mutations in *SPG20* cause a complicated form of HSP known as Troyer syndrome (*7*, *15*). In mouse models, conventional *Spg20* knockout (KO) leads to cognitive and motor deficits, and disruption of Spg20-mediated lipophagy causes neuronal LD accumulation (*7*, *12*), suggesting that proper regulation of LD turnover is essential for neuronal function. Yet, how LD formation is regulated in neurons, how they are mobilized, and how failed LD mobilization contributes to neuronal dysfunction remain unknown.

A major barrier has been the lack of technology capable of detecting neuronal LDs with sufficient specificity and sensitivity in primary cultured neurons and *in vivo*. Electron microscopy offers ultrastructural resolution, but reliable detection of small neuronal LDs can be limited by sample preparation, and lipophilic dyes can produce non-specific labeling of neutral lipid-rich structures in neuronal compartments, thereby hindering sufficient signal-to-noise ratio. Existing LD biosensors have enabled broad LD visualization (*16*, *17*), but their applications to neurons remain limited. Although recent work with tdTomato-PLIN2 knock-in mice has advanced *in vivo* LD visualization (*18*), techniques capable of resolving the heterogeneity of LD populations with distinct molecular features remain limited. In this study, we developed LipiDew, a genetically encoded pan-LD reporter, and used it to demonstrate that neuronal activity induces LD formation in neurons and that regulated LD degradation is required for neuronal function.

To develop a genetically encoded probe that targets LDs with high selectivity and sensitivity, we tested protein domains reported to target LD surfaces from the endoplasmic reticulum (ER) or cytoplasm (*19*). The candidates included ER-derived membrane anchors that access LDs from the ER and cytosolic amphipathic helices that associate directly with the LD surface (*20*). An optimized construct, derived from a liver-enriched LD-associated protein HSD17B13, consists of an N-terminal hydrophobic domain (aa 1–28) and a short C-terminal helix–turn–helix motif (aa 260–286) (*21*, *22*). This chimeric protein showed selective LD targeting and was termed LipiDew (Fig. 1A and fig. S1). In the absence of LD induction, LipiDew was localized to the ER through its hydrophobic domain (aa 1–28) but rapidly redistributed to nascent LD surface during LD biogenesis induced by oleic acid (OA) supplementation (Fig. 1, B–E). Structural prediction and sequence analysis suggested that the N-terminal region of HSD17B13 (aa 1–28) forms a monotopic transmembrane domain that inserts shallowly into the ER bilayer, whereas the helix–turn–helix motif (aa 260–286) that has amphipathic property stabilizes its association with the LD surface (*22*) (Fig. 1A). Furthermore, LipiDew expression neither induced ectopic LD formation nor disrupted recruitment of an endogenous LD protein PLIN3 in SUM159 cells (fig. S2).

**Fig. 1.**
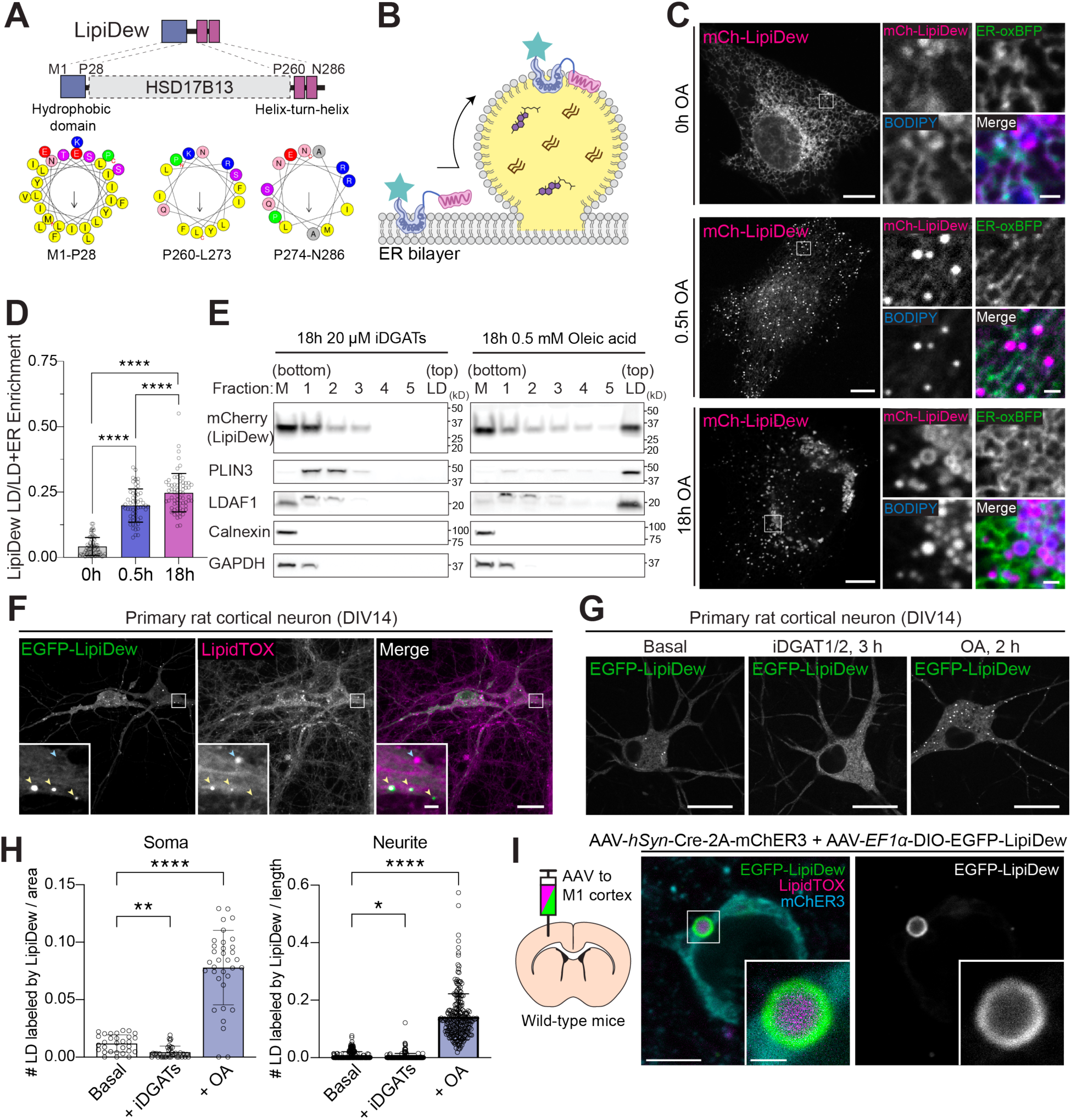
Development of LipiDew, a genetically encoded lipid droplet indicator compatible with brain LD annotation both *in vitro* and *in vivo*. (**A**) Design of LipiDew from the LD-targeting regions of HSD17B13 (Top panel). LipiDew contains the N-terminal hydrophobic segment and C-terminal helix–turn–helix motif of HSD17B13. Helical wheel projections show the predicted α-helical organization of each segment (Bottom panel). (**B**) Schematic showing proposed targeting mechanism of LipiDew to the LD surface. (**C**) Live confocal imaging of SUM159 cells expressing mCherry-LipiDew and ER-oxBFP. Cells were untreated or treated with 0.5 mM OA for the indicated times. LDs were stained with BODIPY493/503. Scale bars: full, 10 µm; insets, 1 µm. (**D**) Quantification of LipiDew enrichment on LDs relative to total signal on the ER and LDs from (C). Mean ± s.d.; *n* = 67, 57, and 64 cells for 0, 0.5, and 18 h OA, respectively, from three independent experiments. Statistical significance was determined by Brown–Forsythe and Welch ANOVA followed by Dunnett’s T3 multiple-comparisons test. (**E**) Subcellular fractionation of SUM159 cells expressing mCherry-LipiDew after 18 h treatment with 20 μM iDGAT1/2 or 0.5 mM OA. Fractions from the bottom to the top of a discontinuous sucrose gradient were analyzed by immunoblotting for LipiDew, PLIN3 (CYTOLD LD marker), LDAF1 (ERTOLD LD marker), Calnexin (ER marker), and GAPDH (Cytosol marker). (**F**) Live confocal imaging of primary rat cortical neurons (DIV14) expressing EGFP-LipiDew and stained with LipidTOX Deep Red to visualize LDs. Yellow arrowheads indicate LipiDew-and LipidTOX-positive puncta; Cyan arrowheads indicate LipidTOX-positive structures lacking detectable LipiDew signal. Scale bars: full, 20 μm; insets, 2 μm. (**G**) Live confocal imaging of primary rat cortical neurons expressing EGFP-LipiDew at DIV14 under basal conditions, after 3 h DGAT1/2 inhibition, or after 2 h OA treatment. Scale bars, 20 μm. (**H**) Quantification of LipiDew-labeled LDs in soma and neurites under the conditions shown in (G) Data are mean ± s.d.; soma, *n* = 30, 35, and 34 cells; neurites, *n* = 214, 216, and 234 neurites for basal, iDGATs, and OA conditions, respectively. Statistical significance was determined by Kruskal–Wallis test followed by Dunn’s multiple-comparisons test comparing each condition with basal. \**p* < 0.05; \*\**p* < 0.01, \*\*\*\**p* < 0.0001. (**I**) *In vivo* labeling of neuronal LDs in mouse M1 cortex after co-delivery of AAV-*hSyn*-Cre-2A-mCherry-ER3 (luminar ER marker) and AAV-*EF1α*-DIO-EGFP-LipiDew. LipiDew-positive structures colocalized with LipidTOX-positive LDs. Scale bars: full, 5 μm; insets, 1 μm.

A criterion we set for a robust LD indicator in neurons was to ensure selective LD annotation without indicator-induced protein aggregation. As shown in Fig. 1F, adeno-associated virus (AAV)-mediated expression of EGFP-LipiDew under the neuronal *Synapsin* promoter selectively labeled LDs in primary rat hippocampal neurons with high specificity, unlike the conventional lipophilic dye LipidTOX, while showing no detectable protein aggregation. In addition, LipiDew labeling remained tightly coupled to cellular triglyceride (TG) content: pharmacological inhibition of DGAT1 and DGAT2, the enzymes catalyzing the final step of TG synthesis (*23–25*), reduced LipiDew-positive LDs, whereas OA supplementation robustly increased LD abundance in both soma and neurites (Fig. 1, G and H). Lastly, AAV-mediated *in vivo* expression enabled robust visualization of LDs in mouse M1 cortical neurons, clearly distinguishable from the ER lumen marker mCherry-ER3 (Fig. 1I), indicating that LipiDew can mark LDs with sufficient specificity in cultures as well as in intact mouse brains.

Although neurons have not been considered LD-rich cells under physiological conditions, disruption of lipolysis and lipophagy causes LD accumulation in neurons (*7*, *13*), suggesting that neuronal LDs undergo active turnover. Because neuronal activity acutely reshapes cellular metabolism and membrane homeostasis, we hypothesized that neuronal activation may modulate LD biogenesis. The ability of LipiDew to selectively visualize neuronal LDs in primarily cultured neurons and mouse brains provided us with a unique opportunity to examine how neuronal activity regulates LD dynamics.

To test whether neuronal activity modifies LD dynamics *in vivo*, we employed a complex rotarod paradigm task (*26*, *27*) in mice expressing LipiDew and mCherry-ER3 in the primary motor cortex (M1) (Fig. 3A and fig. S3). Motor exercise in this task activated the M1 cortex, as indicated by c-Fos induction (fig. S3), and induced LD accumulation (Fig. 2, A–E). LipiDew-positive LDs were increased in both somatic and extrasomatic compartments of motor cortical neurons in exercised mice compared to those in naïve controls, but the degree of LD induction was more prominent in extrasomatic compartments than soma (Fig. 2, B and C). In addition, the volume of somatic LDs was increased while the volume of extrasomatic LDs remained unchanged (Fig. 2, D and E).

**Fig. 2.**
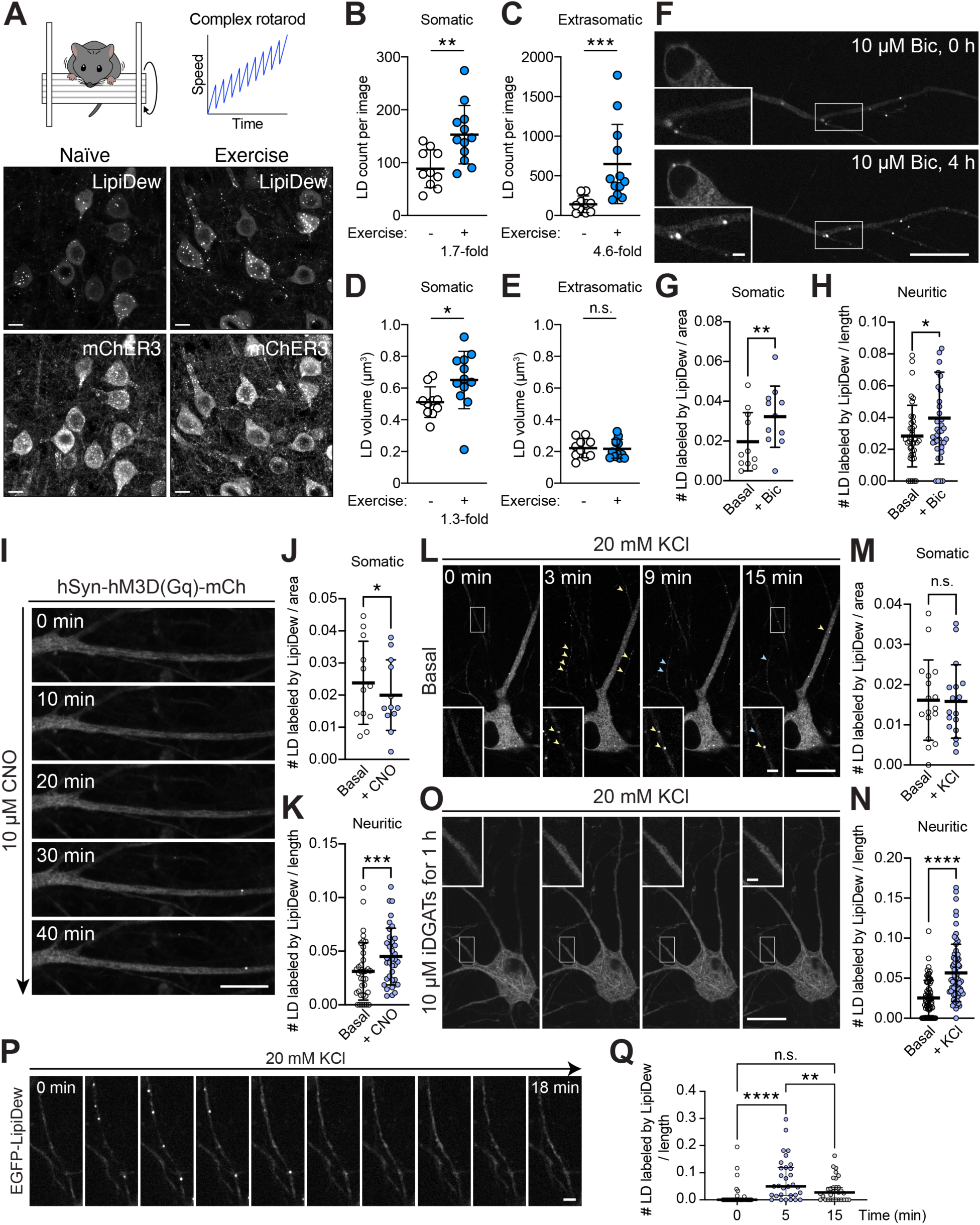
Activity-dependent LD formation in neurons. (**A**) The complex rotarod paradigm. Ten repeats of 5-min dynamic rotarod sessions were used, as illustrated in the graph, with 5-min inter-trial intervals. AAV-injected, but non-trained, naïve littermate mice in home cages were used as control. *In vivo* labeling of neuronal LDs in the mouse M1 cortex after co-delivery of AAV-*hSyn*-Cre-2A-mCherry-ER3 (luminal ER marker) and AAV-*EF1α*-DIO-EGFP-LipiDew from naïve and exercised mice. Scale bars: 10 μm. (**B** to **E**) Quantification of somatic (**B**) and extrasomatic (**C**) LD counts and somatic (**D**) and extrasomatic (**E**) LD volume in naïve and exercised mice. Mean ± s.d., *n* = 10–12 images from *n* = 4 mice per condition, \**p* < 0.05, \*\**p* < 0.01, \*\*\**p* < 0.001, n.s., non-significant, Mann–Whitney tests. (**F**) Live confocal imaging of LipiDew in primary cortical neurons (DIV16) treated with 10 μM bicuculline for 0 or 4 h. Insets show LipiDew-positive puncta in neurites. Scale bars: full, 20 μm; insets, 2 μm. (**G** and **H**) Quantification of LipiDew-labeled LDs in soma (G) and neurites (H) under basal conditions or after bicuculline treatment. Mean ± s.d.; soma, *n* = 11 cells; neurites, *n* = 41 neurites for 0 h and 4 h conditions, respectively. Statistical significance was determined by a two-tailed Wilcoxon matched-pairs signed-rank test. \**p* < 0.05; \*\**p* < 0.01. (**I**) Time-lapse confocal imaging of LipiDew in neurites of primary cortical neuron (DIV14) expressing AAV-*hSyn*-hM3D(Gq)-mCherry before and after 10 μM CNO treatment. Scale bar: 10 μm. (**J** and **K**) Quantification of LipiDew-labeled LDs in soma (J) and neurites (K) under basal conditions or after CNO treatment. Mean ± s.d.; soma, *n* = 12 images per condition; neurites, *n* = 36 neurites per condition. Statistical significance was determined by a two-tailed Wilcoxon matched-pairs signed-rank test. \**p* < 0.05; \*\*\**p* < 0.001. **(L)** Time-lapse confocal imaging of LipiDew in primary cortical neurons (DIV14) before and after 20 mM KCl stimulation. Arrowheads indicate LipiDew-positive puncta appearing or persisting (yellow) and disappearing (blue) during stimulation (yellow). Scale bar: full, 20 μm; insets, 2 μm. (**M** and **N**) Quantification of LipiDew-labeled LDs in soma (M) and neurites (N) under basal conditions or after KCl stimulation. Mean ± s.d.; soma, *n* = 17 images per condition; neurites, *n* = 70 neurites per condition. Statistical significance was determined by a two-tailed Wilcoxon matched-pairs signed-rank test. n.s., not significant; \*\*\*\**p* < 0.0001. (**O**) Time-lapse imaging of LipiDew in primary neurons pretreated with 10 μM DGAT inhibitors for 1 h and stimulated with 20 mM KCl. Scale bars: full, 20 μm; insets, 2 μm. (**P**) Representative time-lapse montage of LipiDew signals in neurites during KCl stimulation (DIV19). Scale bar, 2 μm. (**Q**) LipiDew-labeled LDs were quantified in the same neurites over time following KCl stimulation at 0, 5, and 15 min. LD number was normalized to neurite length. n = 31. Mean ± s.d.; *n* = 31 neurites from 10 neurons. Statistical significance was determined by a Friedman test with Dunn’s multiple comparisons test. n.s., not significant; \*\**p* < 0.01; \*\*\*\**p* < 0.0001.

**Figure 3.**
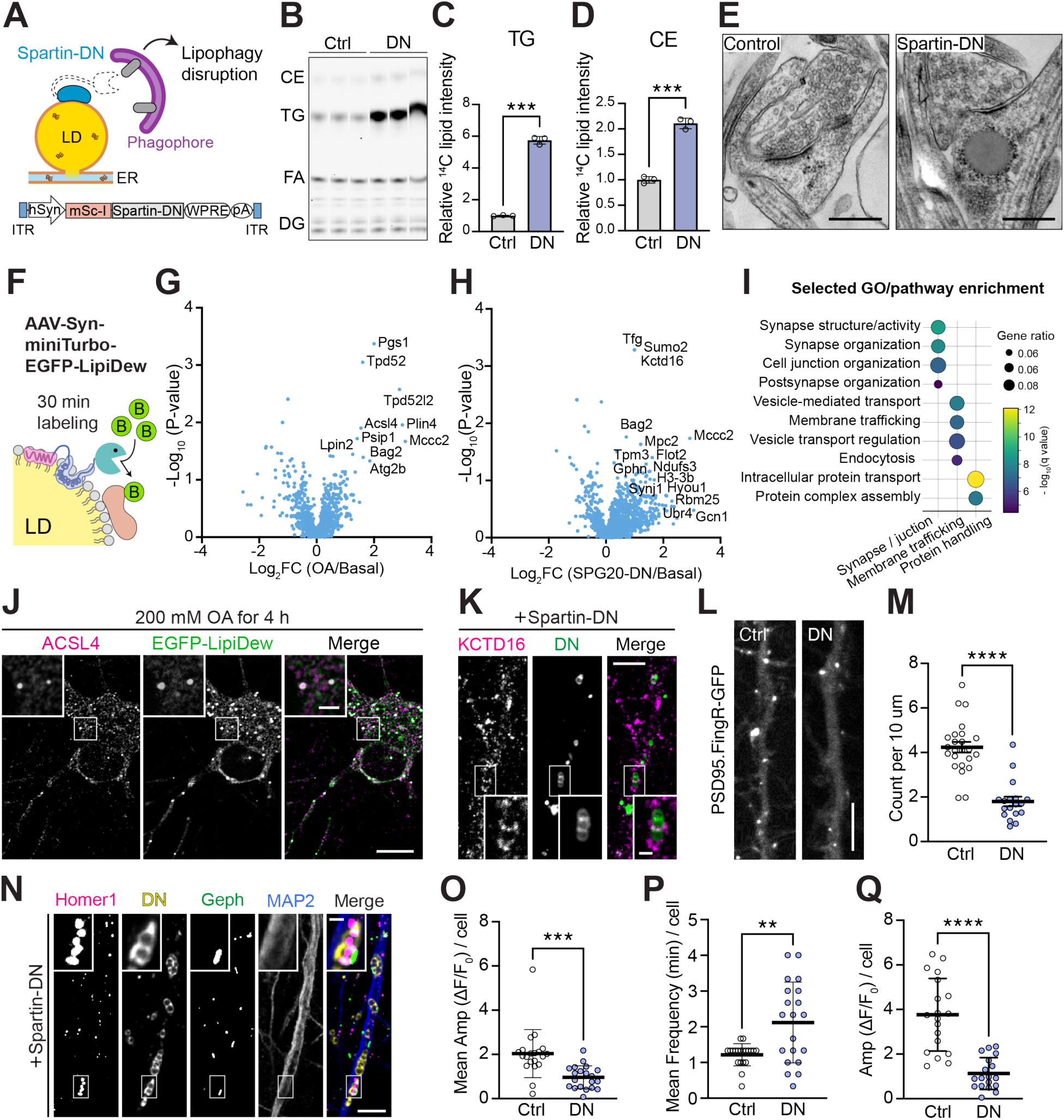
Lipophagy disruption impairs spine integrity and promotes protein mistargeting to lipid droplets. (**A**) Schematic of lipophagy disruption using dominant-negative Spartin (Spartin-DN). (**B**) Thin-layer chromatography analysis of neutral lipids in control and Spartin-DN-expressing primary cortical neurons (DIV16). CE, cholesteryl ester; TG, triglyceride; FA, fatty acid; DAG, diacylglycerol. (**C** and **D**) Quantification of TG (C) and CE (D) levels from (B). Mean ± s.d.; *n* = 3 independent samples per condition. Statistical significance was determined by two-tailed Welch’s t-test. \*\*\**p* < 0.001. (**E**) Electron micrographs of dendritic regions from control and Spartin-DN-expressing primary cortical neurons (DIV19). Scale bars, 500 nm. (**F**) Strategy for proximity labeling of LD-associated proteins using AAV-*hSyn*-miniTurbo-EGFP-LipiDew. (**G** and **H**) Volcano plots showing proteins enriched near LipiDew-labeled LDs after OA treatment (**G**) or Spartin-DN expression (**H**) relative to basal conditions in primary cortical neurons (DIV14). Selected enriched proteins are labeled. (**I**) Selected GO/pathway enrichment analysis of proteins enriched near LDs in Spartin-DN-expressing neurons. (**J**) Confocal images of OA-treated primary cortical neurons (DIV14) expressing EGFP-LipiDew and immunostained for ACSL4. Scale bars, full, 10 μm; insets, 2 μm. (**K**) Confocal images of Spartin-DN-expressing primary cortical neurons (DIV18) immunostained for KCTD16. Scale bars, full, 5 μm; insets, 1 μm. (**L**) Representative dendritic images of PSD95.FingR-GFP in control and Spartin-DN-expressing primary cortical neurons (DIV14). Scale bar, 5 μm. (**M**) Quantification of PSD95.FingR-GFP puncta along dendrites in (L). Data are mean ± s.d.; *n* = 24 control and 18 Spartin-DN dendritic segments. Statistical significance was determined by two-tailed Mann–Whitney test. \*\*\*\**p* < 0.0001. (**N**) Confocal images of Spartin-DN-expressing primary cortical neurons (DIV18) immunostained for Homer1, gephyrin, and MAP2. Scale bars, full, 5 μm; insets, 1 μm (**O** and **P**) Quantification of spontaneous calcium transient amplitude (**O**) and frequency (**P**) in control and Spartin-DN-expressing primary cortical neurons (DIV18–19) expressing jGCaMP7b. Mean ± s.d.; *n* = 20 control and 20 Spartin-DN cells. Statistical significance was determined by two-tailed Welch’s t-test. \*\**p* < 0.01; \*\*\**p* < 0.001.(**Q**) Quantification of glutamate-evoked calcium transient amplitude in control and Spartin-DN-expressing primary cortical neurons (DIV14) in the presence of TTX. Mean ± s.d.; *n* = 19 control and 18 Spartin-DN cells. Statistical significance was determined by two-tailed Welch’s t-test. \*\*\*\**p* < 0.0001.

To characterize the kinetics of activity-induced neuronal LD formation, we expressed LipiDew in primary cortical neurons and assessed LD dynamics with three paradigms of neuronal activation. First, treatment of bicuculline (a GABA_A_ receptor antagonist) increased LD formation in neurites over four hours (Fig. 2, F–H). To test LD dynamics during acute neuronal activation, we next expressed hM3Dq-mCherry, an excitatory chemogenetic receptor activated by clozapine-N-oxide (CNO), and found that CNO treatment increased LD formation in neurites (Fig. 2, I–K). Lastly, acute depolarization via KCl treatment rapidly triggered LD formation along neurites within minutes of the stimulation (Fig. 2, L–N). This response was abolished by inhibition of DGAT1 and DGAT2-mediated triglyceride synthesis, indicating that activity-dependent LD formation requires *de novo* neutral lipid synthesis (Fig. 2O). Whereas OA loading induced LD formation in both soma and neurites (Fig. 1H), activity-dependent LD accumulation occurred predominantly in neurites rather than soma across all stimulation paradigms, consistent with our *in vivo* data based on motor cortex stimulation (Fig. 2, A–E). These findings suggest a differential regulation and function of LDs in neuronal compartments in response to neuronal activity. More interestingly, LD formation was transient in neurites at the peak of 5–10 min after depolarization, and the newly formed LDs disappeared over 30 min, suggesting active clearance mechanisms of neuronal LDs (Fig. 2, P and Q). Together, these results suggest that neuronal LD biogenesis is a rapid, transient, and compartment-involved response to neuronal activity.

The transient nature of activity-dependent LD formation suggested that neuronal LDs are actively turned over. We therefore tested whether impaired LD clearance affects neuronal function. Spartin/SPG20 is a lipophagy receptor whose disruption causes robust accumulation of neutral lipids and LDs in cortical neurons (*7*), and whose mutations have been reported to cause Troyer Syndrome, a form of HSP associated with motor function deficits in patients (*28*). To determine how impaired lipophagic LD clearance affects neuronal functions, we disrupted lipophagic LD degradation using a dominant-negative form of spartin (Spartin-DN). Spartin-DN retains the LD-targeting region of spartin but cannot engage the autophagic machinery, thereby preventing LD delivery to lysosomes without altering TG synthesis (*7*) (Fig. 3A). Primary cortical neurons expressing Spartin-DN showed a substantial increase in TG and cholesteryl esters (Fig. 3, B–D). Along with this neutral lipid accumulation, lipophagic disruption resulted in abnormal electron-dense deposits of LDs in dendritic spines and shafts, implicating postsynaptic regions as a target site of lipid dysregulation (Fig. 3E and fig. S4).

To further characterize how impaired lipophagic LD clearance reshapes the landscape of LD-associated proteomes, we expressed LipiDew fused with miniTurbo, an engineered proximity-labeling enzyme that biotinylates nearby proteins (*29*), in cortical neurons and performed proteomic profiling of neuronal LDs (Fig. 3F, fig. S5). We compared proteins enriched on LDs under basal conditions with those enriched in OA-treated or Spartin-DN-expressing neurons. Gene Ontology analysis revealed that OA-induced neuronal LDs were enriched for canonical LD proteins involved in lipid synthesis and storage, including PLIN4 and ACSL4 (*30*, *31*) (Fig. 3, F and G; fig. S5). By contrast, LDs that were accumulated upon lipophagy disruption via Spartin-DN were enriched for non-canonical LD-associated proteins involved in synaptic organization, membrane trafficking, and regulation of protein-complex assembly, including synapse-regulatory and chaperone-associated factors such as KCTD16 and BAG2 (*32*, *33*) (Fig. 3, H, I, and K; fig. S5).

Disruption of lipophagy by Spartin-DN also compromised synaptic integrity. Neurons expressing Spartin-DN showed a significant reduction in dendritic spines, visualized by PSD95.FingR, an anti-PSD-95 intrabody (*34*) (Fig. 3, L and M), indicating that impaired LD turnover directly affects postsynaptic structure. Consistent with this synaptic phenotype, immunostaining further revealed that dendritic LDs accumulated in proximity to both excitatory and inhibitory postsynaptic markers, Homer1 and gephyrin, respectively (Fig. 3N). Together, these results suggest that failure to clear activity-induced LDs disrupts dendritic synaptic architecture and is associated with aberrant LD accumulation at postsynaptic sites.

We next asked whether these synaptic changes were accompanied by altered neuronal activity by performing calcium imaging in primary cortical neurons expressing jGCaMP7b, a genetically encoded calcium indicator (*35*). We found that neurons expressing Spartin-DN had reduced calcium transient amplitudes and increased event frequency compared with controls in basal conditions (Fig. 3, O and P; fig. S6A). In addition, glutamate-evoked calcium responses in the presence of tetrodotoxin were also markedly blunted (Fig. 3Q; fig. S6B), suggesting impaired postsynaptic responsiveness. Collectively, these comprehensive findings suggest that defective LD clearance reshapes the molecular identity of neuronal LDs, compromises synaptic integrity, and disrupts neuronal activity.

Lastly, we asked whether neuron-specific impairment of lipophagic LDs clearance is sufficient to produce functional defects *in vivo*. We expressed Spartin-DN pan-neuronally under the *Synapsin* promoter using systemic AAV-PHP.eB delivery and performed grip strength tests and open field tests (OFT) (*36*, *37*) (Fig. 4A). Long-term neuronal expression of Spartin-DN significantly reduced grip strength performance (Fig. 4, B and C), without significant changes in general locomotion and anxiety-like behavior measured by OFT (fig. S7, A–D), suggesting that neuron-specific disruption of spartin-dependent lipophagy is sufficient to impair motor functions. To corroborate this finding with an independent genetic approach, we generated a floxed Spg20 mouse line and crossed it with Syn1-Cre mice to selectively delete Spg20 expression in neurons (Fig. 4, D and E). Neuron-specific Spg20 KO mice also showed reduced grip strength performance compared with control littermates (Fig. 4, F and G), again without significant changes in locomotion or anxiety-like behavior in OFT (fig. S7, E–H). Together, these approaches with the dominant-negative form and neuron-specific knockout of Spartin/Spg20 provide *in vivo* evidence that neuronal lipophagy is required to maintain motor performance and that its selective disruption is sufficient to impair motor function.

**Fig. 4.**
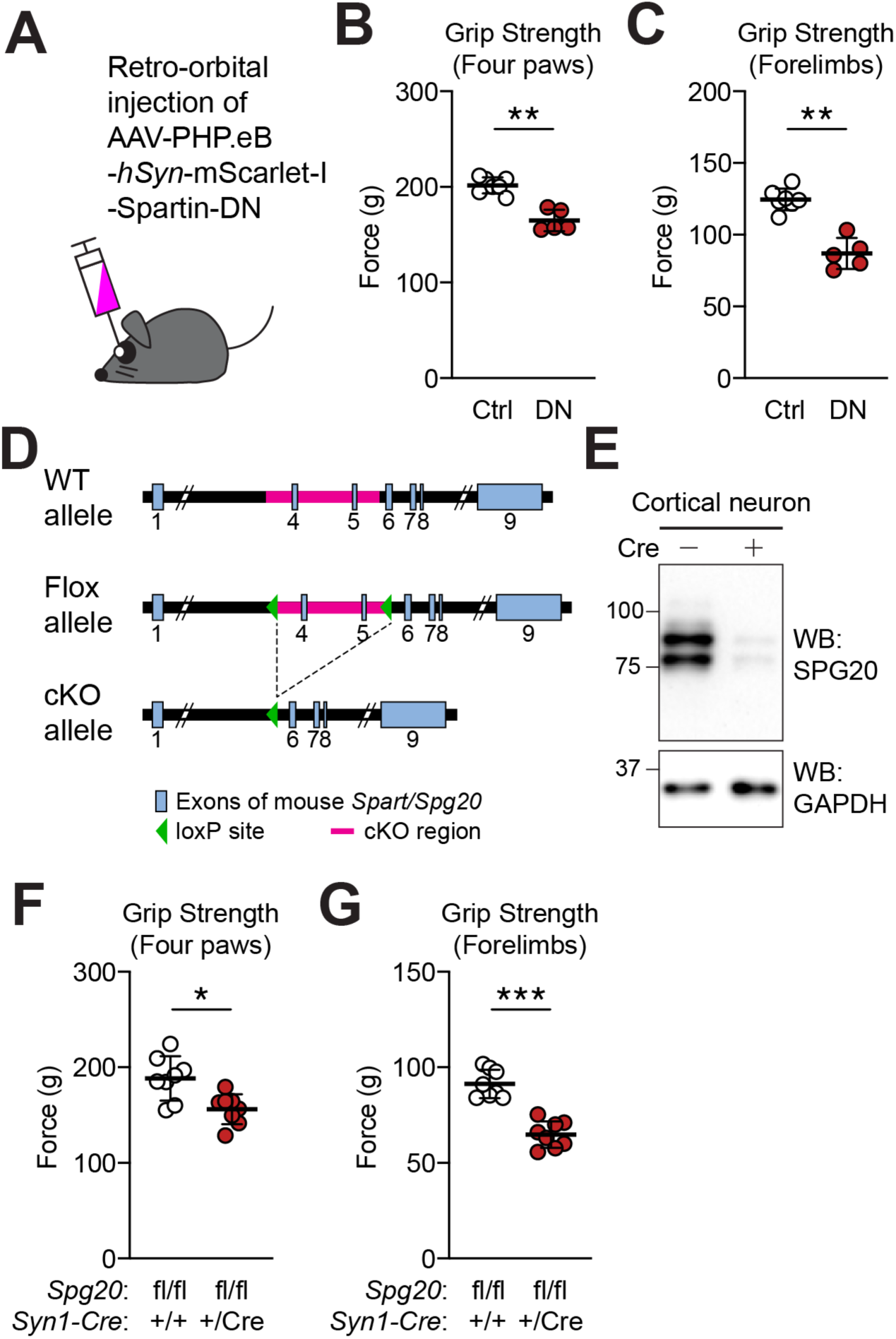
Neuronal lipophagy disruption impairs motor functions. (**A**) Schematic of systemic AAV-PHP.eB delivery by retro-orbital injection to express *hSyn*-mScarlet-I-Spartin-DN in neurons. (**B** and **C**) Grip strength in control and Spartin-DN-expressing mice, measured using four paws (**B**) or forelimbs (**C**). Mean ± s.d.; *n* = 7 control and 5 Spartin-DN mice. Statistical significance was determined by Mann–Whitney tests (\*\**p* < 0.01). (**D**) Schematic of the mouse *Spg20* wild-type, floxed, and conditional knockout alleles. LoxP sites flank the targeted region deleted by Cre recombination. (**E**) Immunoblot analysis of SPG20 in primary cortical neurons (DIV14) from *Spg20* loxP-containing mice treated with 1 μM FUdR/uridine at DIV5 to reduce glial cell growth and transduced with AAV-*hSyn*-Cre at DIV6. GAPDH was used as a loading control. (**F** and **G**) Grip strength in control and neuron-specific *Spg20* knockout mice, measured using four paws (**F**) or forelimbs (**G**). Mean ± s.d.; *n* = 8 mice per genotype. Statistical significance was determined by Mann–Whitney tests (\**p* < 0.05, \*\*\**p* < 0.001).

In this study, we identify neuronal LDs as dynamic organelles that couple neuronal activity to lipid regulation and, in turn, neuronal functions. By combining neuronal LD labeling with neuron-specific perturbations of Spartin/Spg20, we show that neuronal LDs are actively formed and cleared to support neuronal function. Together with recent evidence that neuronal TGs provide a fatty-acid fuel reserve for synaptic function (*14*, *38*), our findings suggest that LD biogenesis and turnover may support both lipid buffering and metabolic supply in active neuronal compartments.

Neuronal activation rapidly and transiently induces LD formation predominantly in neurites, suggesting that LD biogenesis provides a local, transient buffering mechanism to accommodate activity-associated changes in lipid metabolism and membrane demand. This neuron-intrinsic buffering mechanism likely operates in parallel with neuron–glia lipid coupling, as studies in *Drosophila* and mammalian systems have shown that neuronal oxidative or activity-associated stress promotes lipid production and transfer to glia, where these lipids are sequestered in LDs for protection against lipotoxicity (*39*, *40*). Which specific lipid species and upstream signaling pathways initiate this activity-dependent LD biogenesis remains to be defined. When neuronal lipophagy is impaired, neutral lipids accumulate in dendritic compartments, and dendritic LDs aberrantly recruit various proteins, including postsynaptic proteins. Whether these LD-associated proteins directly drive synaptic dysfunction or reflect broader remodeling of dendritic proteostasis remains an important question. Nevertheless, neuron-targeted Spartin-DN expression and neuron-specific Spg20 deletion show that impaired neuronal lipophagy is sufficient to disrupt motor performance. Thus, consistent with the proposed role of LDs as protein-sequestering depots (*41*), our findings suggest that defective LD turnover may convert a physiological lipid-buffering organelle into a pathological lipid–protein sink that disrupts postsynaptic structure, intracellular calcium dynamics, and behavioral outcomes. Together, these findings establish activity-dependent LD formation and lipophagic clearance as a regulated lipid-homeostatic cycle essential for neuronal physiology.

## Acknowledgments

We thank N. Bellono and P. Villar for thoughtful discussions and experimental support, P. E. Castillo for insightful comments and discussions, P. De Camilli, Y. Wu, M. Ericsson for electron microscopy troubleshooting and consultation, and T. Lambert for consultation on imaging analysis. Electron microscopy imaging, consultation, and/or services were performed at the Harvard Medical School Electron Microscopy Facility. Additional reagent gifts are acknowledged in the Materials and Methods.

## Funding

This study was supported by the following funding source.

National Institutes of Health 5R35GM156863 (JC)

Chan Zuckerberg Initiative: 2024-338564 (JC and JP)

Esther A. & Joseph Klingenstein Fund, the Simons Foundation (JC)

The Charles H. Hood Foundation (JC)

Star-Friedman Challenge for Promising Scientific Research (JC)

National Institutes of Health (NIH) R01AG089566 (JP)

National Institutes of Health (NIH) R21AG068423 (JP)

Alzheimer’s Association Research Grant 23AARG-1026776 (JP)

NARSAD Young Investigator Grant (JP)

National Institutes of Health 5R35GM156406 (JAP)

## Author contributions

Conceptualization: EK, JP, JC

Methodology: EK, AS, AL, HH, JAP, WI, JP, JC

Investigation: EK, AS, AL, HH, JAP, WI, JP, JC

Visualization: EK, JP, JC

Funding acquisition: SPG, JP, JC

Project administration: JP, JC

Supervision: JP, JC

Writing – original draft: JP, JC

Writing – review & editing: EK, JP, JC

## Competing interests

Authors declare that they have no competing interests.

## Data, code, and materials availability

All data are available in the main text or the supplementary materials. Further information and requests for resources and reagents should be directed to and will be fulfilled by J.C. or J.P. Requests will be handled according to the policies of Harvard University or Wayne State University regarding MTA and related matters.

## Notes

### Competing Interest Statement

The authors have declared no competing interest.

