## Supplemental Materials for "Activity-dependent lipid droplet biogenesis and turnover regulate synaptic integrity"

**The PDF file includes:**

Materials and Methods

Figs. S1 to S7

### Materials and Methods

#### Cell culture

SUM159 breast cancer cells were obtained from the laboratory of Tomas Kirchhausen (Harvard Medical School) and were maintained in DMEM/F-12 GlutaMAX (Thermo Fisher Scientific) supplemented with 5 µg/ml insulin (Millipore Sigma), 1 µg/ml hydrocortisone (Millipore Sigma), 5% FBS (Thermo Fisher Scientific), 50 µg/ml streptomycin, and 50 U/ml penicillin (Thermo Fisher Scientific). SUM159 cells were plated on 35-mm glass-bottom dishes (MatTek Corp). Where noted, cells were incubated with media containing 0.5 mM oleic acid complexed with essentially fatty acid-free BSA.

Rat embryonic day 18 (E18) cortical or hippocampal tissues, purchased from TransnetYX (Cat #SDECX or #SDEHP), were washed twice with 10 mL of pre-warmed Hibernate-E medium (Thermo Fisher Scientific, #A1247601) before enzymatic dissociation. The enzyme solution, consisting of 3 mg Papain (Worthington Biochemical, #LS003120) and 5 mM L-Cysteine (Millipore Sigma, #C7352) in 6 mL Hibernate-E supplemented with 6 µL Deoxyribonuclease I (Millipore Sigma, #D4527-200KU), was activated at 37°C for 30 minutes and syringe-filtered (0.22 µm) onto the tissue. Tissue was incubated in the enzyme solution for 15 minutes at 37°C, then washed twice with inactivation medium (10% FBS, 1% Penicillin-Streptomycin in Neurobasal). Following mechanical dissociation in 1 mL Hibernate-A, the suspension was centrifuged at 200 × *g* for 2 minutes. The pellet was resuspended in plating medium containing 5% FBS, B27 (Thermo Fisher Scientific, #17504044), GlutaMAX (Thermo Fisher Scientific, #35050061), 0.5% Penicillin Streptomycin (Thermo Fisher Scientific, #15140122) in Neurobasal, and $2 \times 10^5$ cells were plated on PDL-coated (Millipore Sigma, #P7886) 35-mm glass-bottom Fluoro Dishes (WPI, #FD35) with 3 ml of plating media. Three to four hours after the initial plating, 80% of the media was exchanged with neuronal media containing B27, GlutaMAX, and 0.5% Penicillin-Streptomycin in Neurobasal. Every four days after the initial plating, 50% of the media was exchanged with neuronal media.

#### Animals

C57BL/6J wild-type mice (Stock #000664) and Syn1-Cre mice (Stock #003966) were obtained from The Jackson Laboratory. Spg20 floxed mice were generated in this study. Mice were handled and treated in accordance with NIH guidelines and protocols approved by the Institutional Animal Care and Use Committees of Harvard University and Wayne State University. Mice were housed in ventilated cages on a 12-h light/dark cycle with *ad libitum* access to standard laboratory chow and water. Male mice were used for histological analyses, and both male and female mice were used for behavioral tests. Mice used in this study were 2–8 months of age.

#### Generation of Spg20 floxed mice

Spg20 floxed mice were generated by Taconic-Cyagen on a C57BL/6J background using CRISPR/Cas9-mediated genome editing. The targeting strategy inserted loxP sites flanking exons 4–5 of the mouse *Spg20/Spart* gene, with the 5′ loxP site in intron 3 and the 3′ loxP site in intron 5. Cre-mediated deletion of this region is predicted to remove an approximately 3.0-kb conditional knockout region and generate a frameshift loss-of-function allele.

For mouse generation, *Cas9* mRNA, a donor vector containing the loxP sites, and two Spg20-targeting gRNAs were co-injected into fertilized mouse eggs. Founder animals were screened by PCR, followed by sequence analysis, and then bred to wild-type mice to confirm germline transmission. F1 mice carrying the floxed Spg20 allele were used to establish the colony.

For routine genotyping, genomic DNA was analyzed by PCR using primers flanking the 5′ loxP insertion site: F5, 5′-CGTTTGCTCACTCAGTAGAGTATG-3′, and R3, 5′-GCAAGATGAGAATGTGCTCAAAGG-3′. This reaction amplified a 204-bp product from the floxed allele and a 135-bp product from the wild-type allele. Cre-mediated recombination was detected using F5 and R4, 5′-CAGGTTCATCGAACACCAGAAGA-3′, which amplified a 232-bp recombined allele product.

**Special reagents and antibodies**

Janelia Fluor dyes with HaloTag ligands (JF549 and JF646) (Grimm et al., 2015) were kind gifts from Luke Lavis (Janelia Research Campus). BODIPY 493/503 (Cat. #D3922) and HCS LipidTOX Deep Red Neutral Lipid Stain (Cat. #H34477) were purchased from Thermo Fisher Scientific. Oleic acid (Cat. #O1383) was purchased from Millipore Sigma. [1-$^{14}$C] oleic acid (Cat. #ARC0297-50) was purchased from American Radiolabeled Chemicals. A 10 mM oleic acid stock solution was prepared in PBS containing 3 mM fatty acid-free BSA (Millipore Sigma, Cat. #A6003). The solution was incubated in a shaking incubator at 37°C for 1 hour to fully dissolve oleic acid in BSA-PBS, then filtered and stored at −20°C.

Where indicated, the following reagents were used: iDGAT1/2 (a gift from the Farese and Walther laboratory), potassium chloride (Millipore Sigma, Cat. #P5405), bicuculline (Tocris Bioscience, Cat. #0130), clozapine N-oxide dihydrochloride (Tocris Bioscience, Cat. #6329), and tetrodotoxin citrate (Alomone Labs, Cat. #T-550).

Primary antibodies used in this study were as follows: rabbit polyclonal anti-SPG20/Spartin (Proteintech, Cat. #13791-1-AP), rabbit polyclonal anti-mCherry antibody for detection of mScarlet-I (Abcam, Cat. #ab167453), rabbit monoclonal anti-GAPDH (Cell Signaling Technology, Cat. #2118), mouse monoclonal anti-Calnexin (Santa Cruz Biotechnology, Cat. #sc-46669), guinea pig polyclonal anti-PLIN3/TIP47 (Fitzgerald, Cat. #20R-TP001), rabbit polyclonal anti-LDAF1 (Chung et al., 2019), chicken polyclonal anti-MAP2 (Synaptic Systems, Cat. #188006), guinea pig polyclonal anti-Homer1(SYSY Cat # 160004), rabbit polyclonal anti-ACSL4 (Proteintech, Cat. #22401-1-AP), rabbit polyclonal anti-KCTD16 (Thermo Fisher Scientific, Cat. #PA5-62146), mouse monoclonal anti-Gephyrin (Synaptic Systems, Cat. #147011), and guinea pig monoclonal anti-c-Fos (Synaptic Systems, Cat. #226308). HRP-conjugated secondary antibodies against mouse and rabbit IgG were purchased from Santa Cruz Biotechnology. Fluorescent secondary antibodies used in this study were purchased from Thermo Fisher Scientific and included DyLight-conjugated goat anti-chicken IgY (H+L) cross-adsorbed secondary antibody, Alexa Fluor 488-conjugated donkey anti-rat IgG (H+L) highly cross-adsorbed secondary antibody, Alexa Fluor 594-conjugated goat anti-rabbit IgG (H+L) highly cross-adsorbed secondary antibody, and Alexa Fluor 647-conjugated goat anti-guinea pig IgG (H+L) highly cross-adsorbed secondary antibody.

### Plasmid construction

For plasmid construction, DNA fragments were synthesized as gBlocks gene fragments (Integrated DNA Technologies) or amplified by PCR using PfuUltra II Fusion HotStart DNA Polymerase (Agilent Technologies), followed by standard restriction enzyme digestion (New England Biolabs). Digested insert and vector fragments were purified by gel extraction using the NucleoSpin Gel and PCR Clean-up Kit (MACHEREY-NAGEL) and ligated using T4 DNA ligase (New England Biolabs). Ligated plasmid constructs were introduced into DH5α competent bacterial cells (Thermo Fisher Scientific) by heat-shock transformation and propagated.

AAV-EF1A-PSD95.FingR-eGFP-CCR5TC was a gift from Xue Han (*1*) (Addgene plasmid #125691; RRID:Addgene_125691). pGP-AAV-syn-jGCaMP7b-WPRE was a gift from Douglas Kim and the GENIE Project (*2*) (Addgene plasmid #104489; RRID:Addgene_104489). pAAV-hSyn-hM3D(Gq)-mCherry was a gift from Bryan Roth (Addgene plasmid #50474; RRID:Addgene_50474). ERmxBFP was a gift from Erik Snapp (Addgene plasmid # 68126 ; RRID:Addgene_68126).

### Lipid droplet fractionation

SUM159 cells were transfected and treated as described above. Following treatment, cells were washed twice with ice-cold DPBS, harvested by scraping, and centrifuged at 500 × g for 10 min. The supernatant was discarded, and the cell pellet was resuspended in isotonic homogenization buffer containing 20 mM HEPES/KOH, pH 7.5, 70 mM sucrose, 220 mM mannitol, 1 mM EDTA, 2 mg/ml BSA, and EDTA-free cOmplete Protease Inhibitor Cocktail tablets (Millipore Sigma). The suspension was transferred to a chilled Dounce homogenizer (Millipore Sigma, Cat. #D8938). Cells were homogenized with 10 strokes using a loose-fitting pestle without additional pressure, followed by 100 strokes using a tight-fitting pestle with constant pressure. The homogenate was centrifuged at 1,000 × g for 10 min to pellet nuclei and unbroken cells. The pellet was retained for later analysis, and the post-nuclear supernatant was transferred to a fresh tube. The post-nuclear supernatant, 600 µl, was mixed with 300 µl of 47.2% sucrose buffer to generate a 20% sucrose solution, which was loaded at the bottom of a TLS-55-compatible ultracentrifuge tube. A 5% sucrose solution, 850 µl, followed by 450 µl of 0% sucrose solution, both prepared in hypotonic homogenization buffer containing 20 mM HEPES/KOH, pH 7.5, 1 mM EDTA, 2 mg/ml BSA, and EDTA-free cOmplete Protease Inhibitor Cocktail tablets, were layered above the sample. Gradients were centrifuged in a TLS-55 rotor at 166,000 × g for 3 hours at 4°C with no brake. The top LD fraction was collected using a centrifuge tube slicer (Beckman Coulter, Cat. #303811), followed by collection of five approximately equal-volume fractions, each ~350 µl, beneath the LD fraction until only the membrane pellet remained. The membrane pellet was resuspended in an equivalent volume, 350 µl, of isotonic homogenization buffer. Triton X-100, 10% in isotonic homogenization buffer, was added to each fraction to a final concentration of 1%, and all fractions were solubilized by rotation overnight at 4°C. Protein enrichment in each fraction was determined by immunoblotting.

### Immunoblotting

For immunoblot analyses, proteins were transferred from gels to Immun-Blot polyvinylidene fluoride membranes (Bio-Rad) in 1× Tris/glycine transfer buffer containing 20% methanol for 1 hour at 70 V in a cold room. Membranes were blocked in TBS-T supplemented with 5% non-fat dry milk (Santa Cruz Biotechnology) at room temperature for 20–60 min and then incubated with primary antibodies overnight at 4°C with gentle shaking. Membranes were washed three times in TBS-T for 5 min each and incubated with appropriate HRP-conjugated secondary antibodies (Santa Cruz Biotechnology) for 1 hour at room temperature. Immunoreactive bands were detected by chemiluminescence using SuperSignal West Pico or SuperSignal West Dura reagents (Thermo Fisher Scientific).

**AAV production**

AAVs for focal infection were generated using the AAV-DJ Helper Free system (Cell Biolabs), as previously described (*3*–*5*). Each AAV construct was co-transfected with pAAV-DJ and pHelper into 293FT cells (Invitrogen). Transfected cells were lysed three days after transfection, and AAVs were purified with 1-ml HiTrap heparin high-performance columns (Cytiva) as described (*3*, *4*). The titer of each AAV was determined by semi-quantitative PCR. AAV-PHP.eB for systemic infection was generated by Vector Biolabs.

**AAV injection**

For focal injections, C57BL/6 wild-type mice (2 months of age) were anesthetized with 1.5–3.0% isoflurane and placed in a stereotaxic apparatus (Kopf Instruments). A small hole was made in the skull over the M1 motor cortex based on stereotactic coordinates. Then, 0.7 µl of AAV was injected into the M1 cortex using a glass pipette (tip diameter 7–10 µm) at a rate of 100 nl/min using a syringe pump (Micro2T; World Precision Instruments). We injected each AAV with the following titer per hippocampus: $7.3 \times 10^7$ viral genomes (vg) of AAV-EF1α-DIO-mEGFP-LipiDew, $2.0 \times 10^6$ vg of AAV-EF1α-DIO-EGFP-LipiDew, and $2.0 \times 10^7$ vg of AAV-hSyn-mCherry-ER3. The injection site was standardized across animals using stereotaxic coordinates (ML = +1.50, AP = +1.00, DV = –1.50 and –1.25) from bregma. After the injections, we waited 10 min before retracting the pipette. After allowing time for recovery and protein expression, mice underwent tissue harvest or behavioral paradigms. For systemic infections, intravenous administration of AAV-PHP.eB-*hSyn*-mScarlet-I-Spartin-DN or mScarlet-I alone ($1.0 \times 10^{11}$ vg) was performed by injection into the retro-orbital sinus of wild-type mice (at 3 weeks of age).

**Viral transduction in primary neuronal cultures**

For live imaging of primary cortical neurons, neurons were transduced with AAVs at days *in vitro* (DIV) 5–6. To disrupt Spartin-mediated lipophagy, neurons were transduced with AAV-*hSyn*-mScarlet-I-Spartin-DN. Where noted, neurons were transduced with AAV-*EF1α*-PSD95.FingR-EGFP, AAV-syn-jGCaMP7b, AAV-*hSyn*-Cre-P2A-mTagBFP2, AAV-*EF1α*-DIO-mEGFP-LipiDew, or AAV-*hSyn*-hM3D(Gq)-mCherry for confocal microscopy. For proximity labeling assays, neurons were transduced with either AAV-*EF1α*-DIO-EGFP-miniTurbo-HA-LipiDew or AAV-*EF1α*-DIO-EGFP-miniTurbo-HA, with AAV-*hSyn*-mScarlet-I-Spartin-DN included where indicated.

### Immunocytochemistry

Primary cortical neurons were fixed with 4% paraformaldehyde (EMS, #15710) and 4% sucrose in DPBS for 20 min at room temperature, then quenched with 50 mM $NH_4Cl$ for 3 min. Samples were permeabilized with 0.1% Triton X-100 (Millipore Sigma, #T9284) in DPBS for 10 min, then blocked in 5% bovine serum albumin (BSA; Millipore Sigma, #A6003) in DPBS for 1 hour at room temperature. Samples were then incubated with their respective primary antibody solutions in blocking buffer at 4°C overnight. After three 10-min washes with DPBS, cells were then incubated with their respective secondary antibody solutions in blocking buffer for 1 hour at room temperature. Following three 10-minute washes, cells were mounted on slides using Vectashield antifade mounting medium.

ACSL4 staining: Primary cortical neurons were fixed with 4% paraformaldehyde (EMS, #15710) and 4% sucrose in DPBS for 20 min at room temperature, quenched with 50 mM $NH_4Cl$ for 3 minutes. Samples were permeabilized with 0.05% saponin (Millipore Sigma, #47036) and 0.1% bovine serum albumin (BSA; Millipore Sigma, #A6003) in DPBS for 3 min, then blocked in 0.05% saponin and 5% normal goat serum (Cell Signaling, #5425S) in DPBS for 1 hour at room temperature. Samples were then incubated with their respective primary antibody solutions in blocking buffer for 1 hour at room temperature. After three 5-min washes with DPBS containing 0.05 saponin and 0.2% BSA, cells were then incubated with their respective secondary antibody solutions in blocking buffer for 1 hour at room temperature. Following three additional 5-min washes with DPBS containing 0.05 saponin and 0.2% BSA, cells were rinsed twice with DPBS and imaged in fresh DPBS.

### Proximity-based LD proteomics

Proximity labeling was performed as previously described (*6*), with minor modifications. Before biotin labeling, neurons were treated with either 200 µM oleic acid for 4 h or 20 mM KCl for 30 min. Neurons were then incubated with 100 µM biotin in neuronal medium for 30 min at 37°C. After aspiration of the medium, cells were washed three times with 1 ml of sterile ice-cold ACSF (125 mM NaCl, 5 mM KCl, 2 mM $CaCl_2$, 1 mM MgCl2, 10 mM HEPES, pH 7.4, 25 mM D-glucose in water). Cells were lysed by scraping in 300 µL of freshly prepared lysis buffer (RIPA buffer supplemented with 1x protease inhibitor cocktail and 1 mM PMSF), then incubated on ice for 10 min. Lysates were cleared by centrifugation at 13,000 × *g* for 10 min at 4°C. Lysates from three 35-mm dishes were combined per sample, corresponding to approximately 600,000 neurons. The combined lysate was added to streptavidin-conjugated magnetic beads and rotated for 1 h at 4°C. Beads were washed twice with 1 mL of RIPA buffer (2 min each), once with 1 mL of 1 M KCl (1 min), once with 1 mL of 0.1 M $Na_2CO_3$ (1 minute), once with 1 mL of 2 M urea in 10 mM Tris-HCl pH 8 (1 min), and twice with 1 mL of RIPA buffer (2 min each), with aspiration between each wash. All washes were performed at room temperature. The $Na_2CO_3$ and urea washes were performed without resuspending the beads to minimize protein loss and denaturation. Beads were then washed twice with 200 mM HEPES (pH 8) for subsequent proteomic analysis.

### Protein Digestion

Beads were resuspended in 200 mM HEPES, pH 8.5, and digested at room temperature for 13 h with Lys-C protease at a 100:1 protein-to-protease ratio. Trypsin was then added at a 100:1 ratio, and the reaction was incubated for 6 h at 37°C. Peptides were separated from beads, vacuum centrifuged to near-dryness and desalted via StageTip (*7*).

**Mass Spectrometric Data Acquisition**

Mass spectrometric data were collected on an Orbitrap Astral instrument coupled to a Vanquish Neo UHPLC. Peptides were separated using a 66-min gradient of 5 to 29% acetonitrile in 0.125% formic acid with a flow rate of 350 nL/min. The spray voltage was set at 2800 V. The scan sequence began with an Orbitrap MS1 spectrum with the following parameters: resolution 120,000, scan range 350-1350 Th, automatic gain control (AGC) target "standard," and maximum injection time 50 ms. MS2 spectra were acquired with the following parameters: Astral mass analyzer, AGC target 20,000, maximum injection time 20 ms, isolation window 1.2 Th, normalized collision energy (NCE) 27.00%, and centroid spectrum data type. Dynamic exclusion was set to automatic. The FAIMS compensation voltages (CV) were -35, -45, -50, -55, and -65 V.

**Mass Spectrometric Data Analysis**

Spectra were converted to mzXML via MSconvert (*8*). Database searching included all entries from the rat UniProt reference Database (downloaded: June 2025). The database was concatenated with one composed of all protein sequences for that database in the reversed order. Searches were performed using a 50-ppm precursor ion tolerance for total protein level profiling. The product ion tolerance was set to 0.03 Da. These wide mass tolerance windows were chosen to maximize sensitivity in conjunction with Comet searches and linear discriminant analysis (*9*, *10*). Oxidation of methionine residues (+15.995 Da) was set as a variable modification. Peptide-spectrum matches (PSMs) were adjusted to a 1% false discovery rate (FDR) (*11*, *12*). PSM filtering was performed using a linear discriminant analysis, as described previously (*10*) and then assembled further to a final protein-level FDR of 1% (*12*). Proteins were quantified by spectral counting. For GO analyses, the Metascape webtool version 3.5(*13*) was used with a cutoff for significance ($p < 0.5$) and fold change (greater than 0.5).

**Transmission electron microscopy**

Primary rat cortical neurons were cultured on 12-mm round glass coverslips (VWR, #76355-906) in PDL-coated 35-mm tissue culture dishes and were fixed with 2.5% glutaraldehyde and 2 mM $CaCl_2$ in 0.1 M sodium cacodylate buffer (pH 7.4) (EMS, #15949) for 3 h at room temperature. Samples were transported to the Harvard Medical School Electron Microscopy Facility, where all subsequent processing and imaging were performed.

Briefly, samples were washed four times in 0.1 M sodium cacodylate buffer (pH 7.4) and preincubated with 0.1 M imidazole and 2 mM $CaCl_2$ for 5 min. Samples were then post-fixed in 2% osmium tetroxide ($OsO_4$) in 0.1 M imidazole containing 2 mM $CaCl_2$ for 20 min. After washing with Milli-Q water, samples were incubated in 2% uranyl acetate in Milli-Q water for 1 h. After another round of washes in Milli-Q water, samples were dehydrated through a graded ethanol series for 5 min each: 50%, 70%, 95%, and 100% twice. Samples were embedded in Epon-Araldite resin at 60°C for 48 h, and glass coverslips were removed by immersion in liquid nitrogen. Ultrathin sections (~60 nm) were cut using a Reichert Ultracut-S microtome, mounted onto copper grids, and stained with lead citrate. Sections were examined using a JEOL 1200EX transmission electron microscope equipped with an AMT 2k CCD camera.

**Calcium imaging of primary cultured neurons**

Calcium imaging was performed 10–14 days after neurons were transduced with AAV-syn-jGCaMP7b. Calcium imaging was carried out using the same confocal microscope (Nikon Eclipse Ti2) and stage-top chamber described above, maintained at 37°C, 5% $CO_2$, and 85% humidity. Images were acquired every 300 ms using a 20x CFI Plan Apochromat Lambda D 0.8 NA objective (Nikon). For spontaneous calcium activity, neurons were imaged in culture medium for 3 min. For the glutamate-evoked calcium response, the medium was replaced with ACSF as described above. After a 2-min baseline recording, 1 µM TTX (tetrodotoxin citrate, Alomone Lab, Cat. #T-550) was applied, and neurons were imaged for an additional 2 min. Glutamate was then added to a final concentration of 25 µM (L-glutamic acid, Fischer Scientific, Cat. #ICN19467780) and imaging was continued for 10 min.

**Histology**

Mice were deeply anesthetized and transcardially perfused with 20 ml of PBS and 20 ml of 4% PFA in 0.1 M phosphate buffer (pH 7.4). After post-fixation overnight, 50 µm-thick coronal sections were cut on a vibratome (Leica) at 4°C. To image LipiDew, mCherry-ER3, and LipidTOX signals, slices were washed twice in PBS, stained with PBS containing HCS LipidTOX deep red neutral lipid stain (Thermo Fisher Cat #H34477; 1:3,000 dilution) at RT for 10 min, and mounted on microscope slides with Vectashield antifade mounting medium (Vector Laboratories). Confocal fluorescence images were acquired on a laser scanning confocal microscope (LSM 780; Zeiss) equipped with a 63× (NA 1.4) objective. For c-Fos and DAPI staining, immunostaining of free-floating brain sections was performed as described previously (*4*) with minor modifications. After washing three times with PBS, slices were permeabilized and blocked for 1 h in PBS containing 5% NGS and 0.3% Triton X-100 and incubated overnight at RT with PBS containing 3% NGS, 0.1% Triton X-100, and guinea pig monoclonal anti-c-Fos antibodies (Synaptic Systems, Cat #226308, 1:1,000). Slices were then washed three times with PBS and incubated at RT for 2 h in PBS containing 3% NGS and Alexa Fluor 647-conjugated goat anti-guinea pig IgG (H+L) highly cross-adsorbed secondary antibody (Thermo Fisher Scientific, Cat #A21450, 1:1,000). After washing twice with PBS, slices were stained with 0.2 µg/ml DAPI (Sigma-Aldrich) in PBS at RT for 15 min. After washing twice with PBS, slices were mounted on microscope slides with Vectashield antifade mounting medium (Vector Laboratories) and imaged on a laser scanning confocal microscope (LSM 780; Zeiss) equipped with a 20× (NA 0.8) objective.

**Behavioral paradigms**

All behavioral experiments were conducted with 2- to 8-month-old littermates at the time of testing and randomization of individual animals. All behavioral tests and analyses were performed in a blinded manner to AAV or mouse genotypes. The M1 motor cortex was activated using a complex rotarod paradigm previously described (*14*, *15*), with minor modifications. Mice were placed on a rotarod for 5 min using a complex paradigm in which rotational speed increased from 0 to 50 rpm and alternated between positive and negative acceleration at 30-sec intervals (Supplemental Figure 3B), and the procedure was repeated 10 times with 5-min intertrial intervals (*14*, *15*). Forty-five minutes after the last rotarod session, mice underwent cardiac perfusion for tissue harvest. For grip strength tests, forelimb strength and combined forelimb and hindlimb strength (four paws) were separately assessed as previously reported (*16*, *17*) with minor modifications. Mice were held by the base of the tail and allowed to grip onto either one bar or a set of bars. After being positioned horizontally and parallel to the bar, mice were gently pulled horizontally by the tail until their grip was released. The maximum force at which they gripped the bar was recorded in grams. Grip strength was measured five times with 1-min intertrial intervals, and the average of the middle three scores was plotted.

**Lipid extraction and thin-layer chromatography**

Primary cortical neurons cultured in 35-mm dishes were pulse-labeled with [$^{14}$C]-oleic acid (1 µCi/µmol) for 18 hours at DIV14. Cells were washed twice with ice-cold ACSF and harvested by scraping in 100 µl Milli-Q water. Cell lysates were mixed with 300 µl of $CHCl_3$:MeOH (1:2, v/v), vortexed, mixed with 100 µl of $CHCl_3$, vortexed again, and then mixed with 100 µl of Milli-Q water, vortexed again. After centrifugation at 2000 × *g* for 5 min, the lower organic phase was transferred to a fresh microcentrifuge tube and dried under a stream of nitrogen. Samples were separated by thin-layer chromatography (TLC) using a hexane: diethyl ether: acetic acid solvent system (80:20:1). A TLC plate was exposed to a phosphor imaging cassette for 48 h and imaged using an Azura Sapphire FL Biomolecular Imager. Lipid standards on the TLC plate were subsequently visualized by iodine vapor staining.

**Fluorescence microscopy**

For SUM159 cells, imaging was carried out at 37ºC approximately 24 h after transfection. For fixed samples, cells were washed twice with ice-cold PBS, then incubated with 4% formaldehyde in PBS for 20 min at room temperature. After fixation, cells were washed three times with PBS for 5 min each. Where noted, cells were stained with 0.5 µM BODIPY493/503 and 1 µg/mL Hoechst33342 for 20 min before imaging.

For primarily cultured neurons, live imaging was performed at 37ºC between DIV14 and DIV21, and the specific neuronal age is indicated in the figure legend. Fixed neurons were imaged in 2 ml of DPBS after the immunocytochemistry protocol described above.

For primary cultured neurons, live imaging was performed at 37°C between DIV14 and DIV21, and the specific neuronal age is indicated in the figure legend. Unless otherwise noted, fixed neurons were imaged in 2 ml of ice-cold DPBS after the immunocytochemistry protocol described above.

Spinning disk confocal microscopy was performed using a Nikon Eclipse Ti2 inverted microscope equipped with Perfect Focus and a CSU-W1 spinning disk confocal head (Yokogawa), controlled by NIS-Elements software (Nikon). Live imaging was maintained at 37°C and 5% $CO_2$, with 85% humidity, using a stage-top incubation chamber. Images were acquired with either a 60x CFI Plan Apochromat Lambda D 1.42 NA objective or 100x CFI Plan Apochromat 1.45 NA objective (Nikon) using a Photometrics Prime BSI camera. Cells and fluorophores were excited using 405, 488, 561, and 640 nm lasers, with emission collected through ET455/50, ET525/36, ET605/52, and ET705/72nm bandpass filters, respectively.

For spine analysis, neurons were transduced with AAV-*EF1α*-PSD95.FingR-EGFP and imaged using a 100x CFI Plan Apochromat 1.45 NA objective (Nikon). Z-stacks were acquired using the piezo focusing system over a total depth of 4 μm with a 0.3-μm step size.

**Image processing and quantification**

All acquired images were processed and prepared using Fiji from raw ND2 images generated by NIS Elements. The scaling of images was matched to the objective used.

For cell and soma area quantifications, outlines were manually drawn on the basis of the EGFP-LipiDew channel in primary cortical neuron culture. The area was then quantified using the "Measure" tool in Fiji. For LD number quantification, LDs were identified by the "Find Maxima" tool in Fiji. Neurite length was measured using the NeuronJ plugin in Fiji. All neurites were manually traced, with at least three neurite analyzed per cell. The total number of LDs in the soma or neurite was normalized to soma area or the neurite length, respectively.

For calcium imaging quantification, imaging data were processed and analyzed using Fiji. For each recording, regions of interest (ROIs) encompassing the soma of individual neurons were manually defined. A minimum of 30 cells were measured per condition. ROI mean fluorescence intensity over time was extracted using the Plot Z-axis Profile function (Image > Stacks > Plot Z-axis Profile), which returns the mean ROI intensity for each frame across the recording. Raw fluorescence intensity traces for individual neurons were exported to Excel, with time in the first column and individual cell traces in subsequent columns. For evoked calcium responses, fluorescence values for each cell were normalized as ΔF/F using the average fluorescence from the final minute of TTX recording before glutamate addition as the baseline (F0), according to (F - F0) / F0. Raw traces were further analyzed using a custom script to identify peaks and measure average peak amplitude and event frequency. Peaks were detected independently for each cell using MATLAB's find peaks function and detection parameters, including a minimum peak height and prominence, which were scaled to the mean and standard deviation of each trace. A minimum peak distance was also applied as a parameter to prevent repeated detection of the same event. Peak frequency was calculated as the number of detected events per minute, while mean peak amplitude was calculated as the local amplitude of each peak by measuring the distance of each trough and peak for each trace.

For quantitative analysis of the density of PSD95.FingR puncta, the count and volume of LipiDew puncta, and the percentage of c-Fos-positive cells, each channel image was processed using the surface analysis function in Imaris 10.1 (Oxford Instruments). Thresholding was performed on all surfaces with adjusted background subtraction before segmentation steps. All parameters used for surface analysis were held consistent across imaging data. To quantify PSD95.FingR puncta in cultured neurons, region-growing segmentation was performed based on morphology. In each image, 1–3 dendrites (25–55 μm length) located 40–50μm away from the soma were analyzed for quantification. LipiDew puncta in the M1 cortex underwent region growing segmentation based on intensity. The c-Fos and DAPI signals in the M1 cortex underwent morphological segmentation. Image quantification analyses were performed in a blinded manner to the conditions.

#### Statistical analysis

Results are presented as mean ± standard deviation. The sample size was chosen based on the previous studies (*4*, *5*). Statistical significance between means was calculated using Brown–Forsythe and Welch ANOVA followed by Dunnett's T3 multiple-comparisons test, a two-tailed Wilcoxon matched-pairs signed-rank test, Mann–Whitney tests, Kruskal–Wallis tests, or two-way ANOVA followed by post-hoc Tukey's multiple comparison (Prism 11, GraphPad). Statistical significance is indicated as follows: *$p < 0.05$, **$p < 0.01$, ***$p < 0.001$, ****$p < 0.0001$, and n.s., not significant.

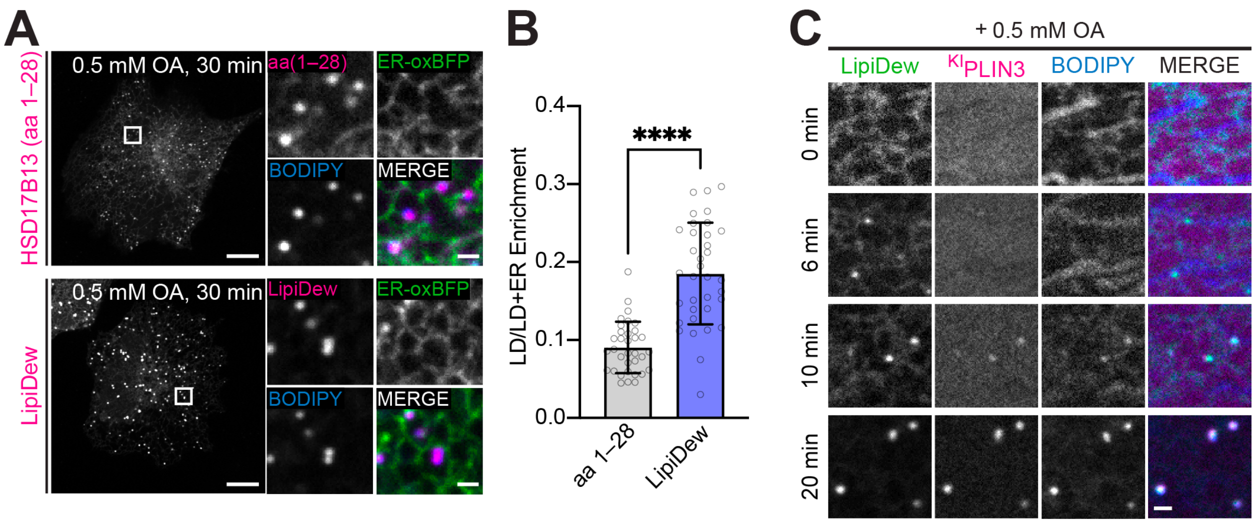


**Fig. S1. LipiDew development from the LD-targeting regions of HSD17B13.**
(**A**) Representative live confocal images of SUM159 cells comparing the N-terminal hydrophobic domain of HSD17B13, HSD17B13 aa 1–28, with LipiDew, which additionally contains C-terminal helix–turn–helix motif. Cells were co-transfected with mCherry-HSD17B (aa 1–28) or mCherry-LipiDew together with ER-oxBFP (an ER luminal marker), stained with BODIPY493/503, and treated with 0.5 mM oleic acid for 30 min. Scale bars: full, 10 µm; insets, 1µm. (**B**) Quantification of LD enrichment relative to the combined LD and ER signal. LipiDew showed significantly greater LD enrichment than HSD17B13 aa 1–28. Mean ± s.d.; individual points represent individual measurements. HSD17B13 aa 1–28, $n = 34$; LipiDew, $n = 35$. ****$p < 0.0001$ by two-tailed Welch's t-test. (**C**) Time-course images showing recruitment of LipiDew during OA-induced LD formation. Cells expressing LipiDew were treated with 0.5 mM OA and stained with BODIPY493/503 and HaloTag was pre-labelled with 100 nM JF646 for 1 h. Scale bar: 1 µm.

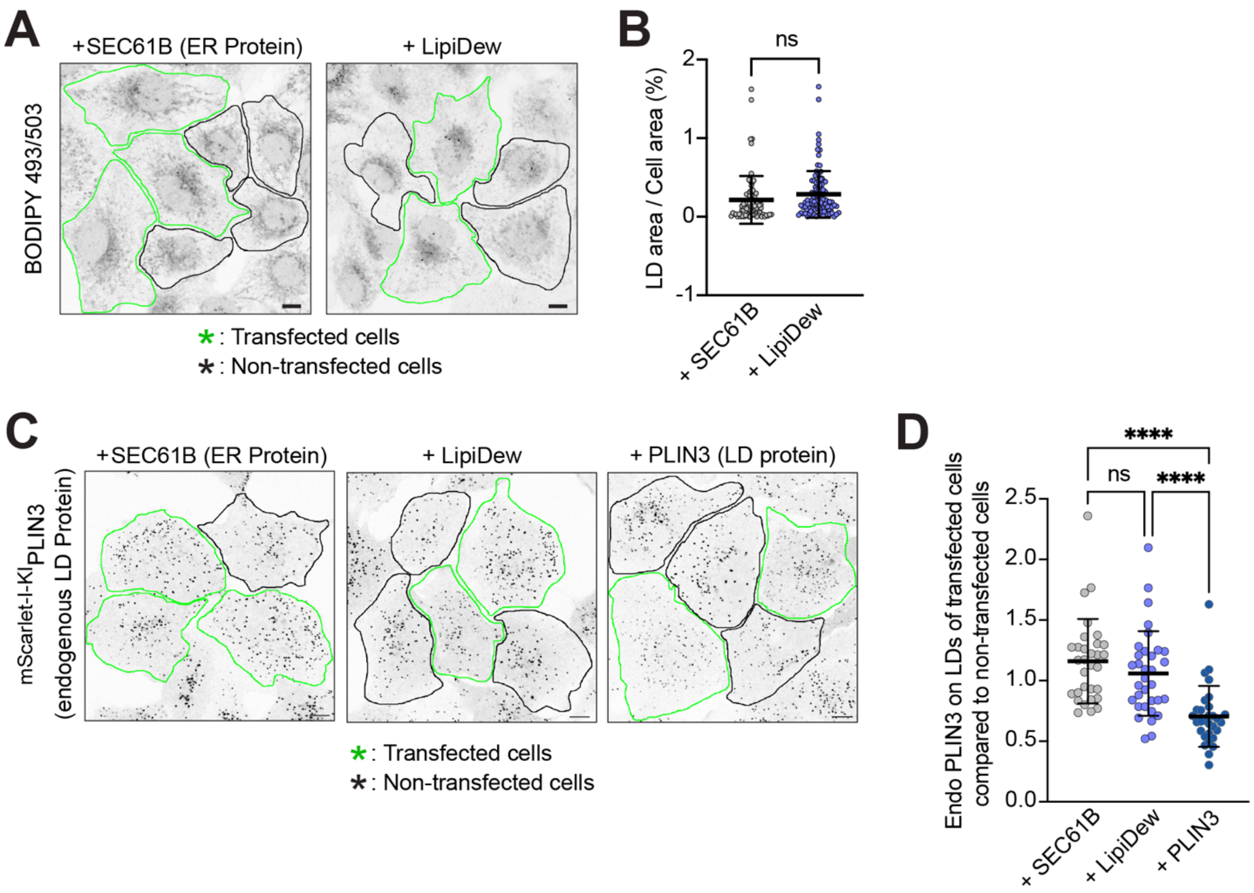


**Fig. S2. LipiDew marks LDs without interfering with endogenous LD properties.**
(**A**) Representative live confocal images of SUM159 WT cells expressing either RFP-Sec61B, an ER marker, or mCherry-LipiDew. LDs were stained with BODIPY493/503. Transfected cells are outlined in green, and non-transfected cells are outlined in black. Scale bars, 10 µm. (**B**) Quantification of LD area normalized to cell area in transfected cells from (**A**). Mean ± s.d.; $n$ = 78 cells for Sec61B and 102 cells for LipiDew, pooled from three independent experiments. n.s., not significant, by two-tailed Welch's unpaired t-test. (**C**) Representative images assessing whether LipiDew expression affects recruitment of endogenous PLIN3 to LDs. SUM159 cells expressing endogenous mScarlet-I-PLIN3 were transfected with HaloTag-Sec61B, HaloTag-LipiDew, or HaloTag-PLIN3 and treated with 0.5 mM OA for 1 h. Cells were then fixed with 4% PFA, and LDs were stained with BODIPY493/503. HaloTag constructs were pre-labeled with 100 nM JF646. Transfected cells are outlined in green, and non-transfected cells are outlined in black. Scale bars, 10 µm. (**D**) Quantification of endogenous PLIN3 enrichment on LDs in transfected cells normalized to neighboring non-transfected cells within the same field of view from (**C**). Mean ± s.d.; $n$ = 31 fields of view for Sec61B, 33 for LipiDew, and 30 for PLIN3, pooled from three independent experiments. n.s., not significant; ****$p < 0.0001$ by Tukey's multiple-comparisons test.

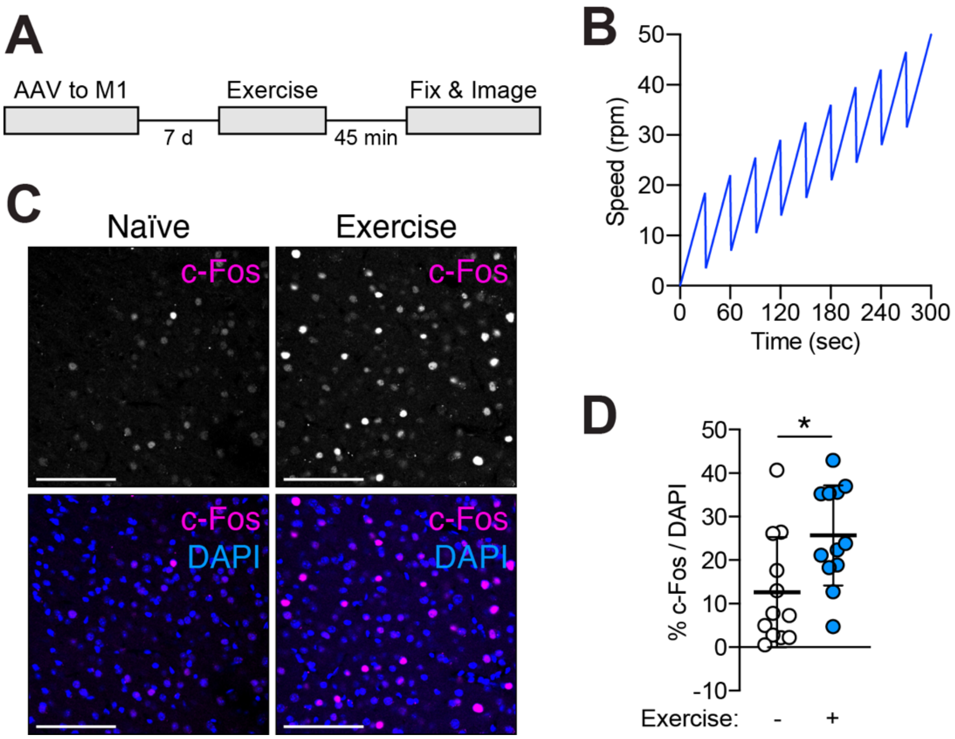


**Fig. S3. Induction of c-Fos expression by a complex rotarod paradigm.**
(**A**) Schematic illustrating experimental procedures. AAV-*hSyn*-Cre-2A-mCherry-ER3 (luminal ER marker) and AAV-*EF1α*-DIO-EGFP-LipiDew were co-injected into the M1 motor cortex of wild-type mice. After seven days, mice were subjected to a complex rotarod paradigm and cardiac perfusion for tissue harvest 45 min after the last rotarod session. (**B**) The complex rotarod paradigm. Ten repeats of 5-min dynamic rotarod sessions were used, as illustrated in the graph, with 5-min inter-trial intervals. AAV-injected, but non-trained, naïve littermate mice in home cages were used as control. (**C**) Increased c-Fos-positive cells in the M1 cortex of trained mice. Scale bars: 100 μm. (**D**) Quantification of the percentage of c-Fos-positive cells against DAPI-positive cells. Mean ± s.d., $n = 12$ images from $n = 4$ mice per condition, $^{*}p < 0.05$, Mann–Whitney tests.

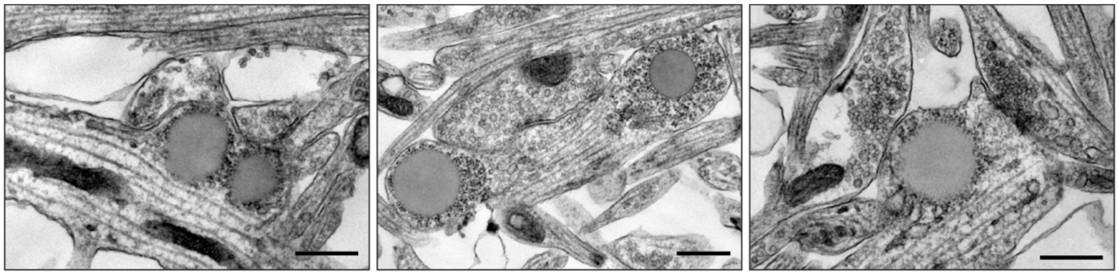

**Fig. S4. Ultrastructure of LD accumulation in Spartin-DN-expressing neurons.** Electron micrographs of synapses from Spartin-DN-expressing primary cortical neurons (DIV19). Scale bars, 500 nm.

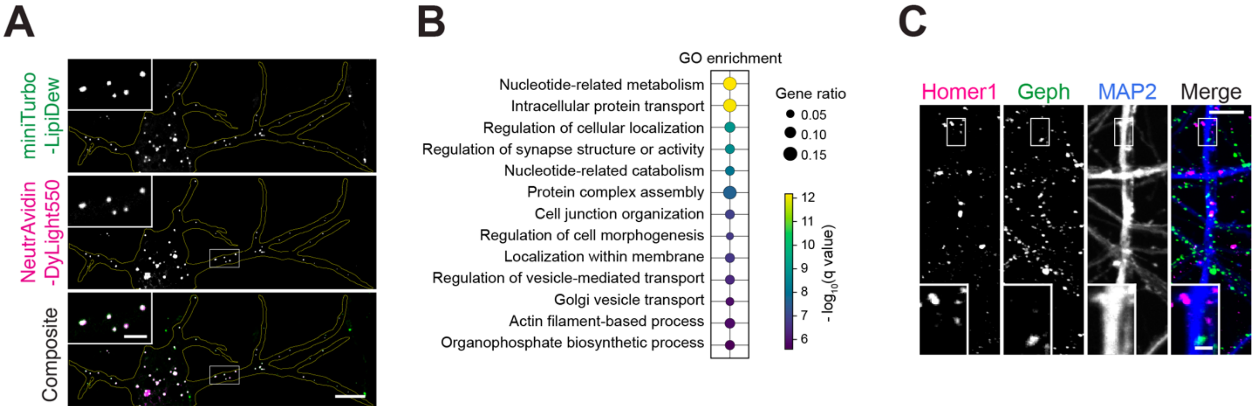


**Fig. S5. LipiDew-TurboID proximity labeling identifies LD-associated proteins in primary neurons.**

(**A**) Proximity-dependent biotinylation of LD-associated proteins in DIV14 primary cortical neurons expressing miniTurbo-LipiDew. Biotinylated proteins were visualized with NeutrAvidin-DyLight550. Neurons were treated with 100 µM OA for 18 h. Scale bars: full, 10 µm; insets, 2 µm. (**B**) Non-curated top GO-term enrichment analysis of proteins enriched near LDs in Spartin-DN-expressing neurons. The corresponding curated GO/pathway analysis is shown in the Fig. 3I. (**C**) Confocal images of control primary cortical neurons (DIV18) immunostained for Homer1, gephyrin, and MAP2. Scale bars, full, 5 µm; insets, 1 µm

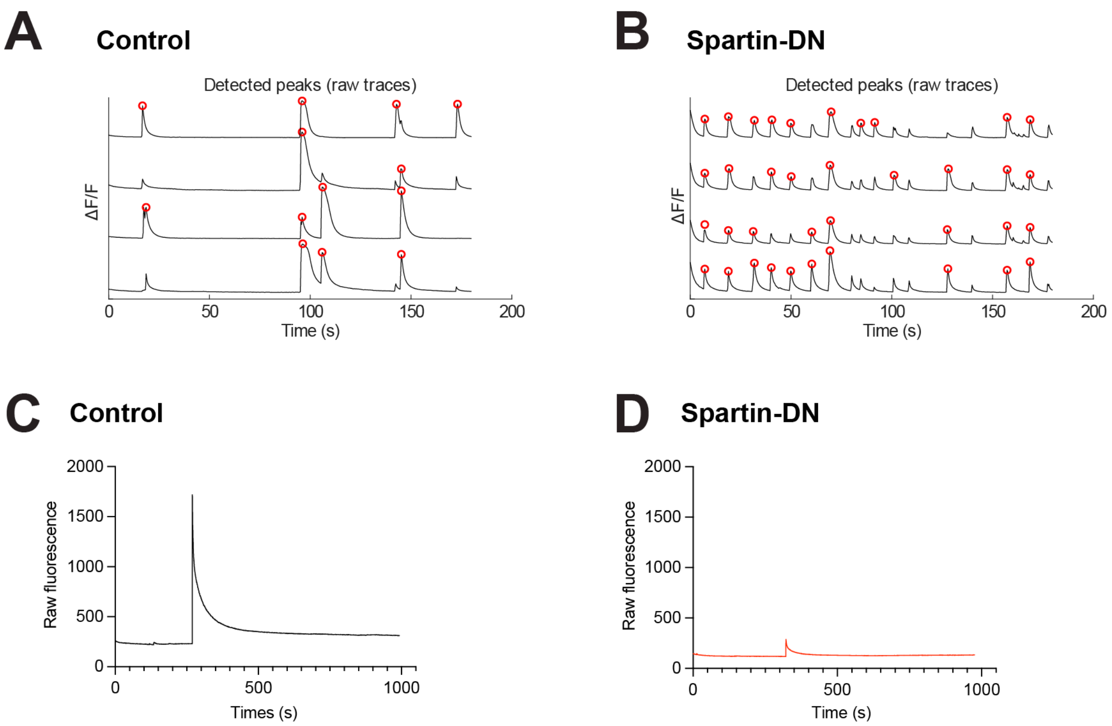


**Fig. S6. Representative calcium-imaging traces from control and Spartin-DN-expressing neurons.**

(**A** and **B**) Representative raw fluorescence traces showing detected spontaneous calcium transients in control (**A**) and Spartin-DN-expressing (**B**) primary cortical neurons expressing jGCaMP7b at DIV18–19. Red circles indicate detected calcium-transient peaks. Quantification is shown in Fig. 3O and 3P. (**C** and **D**) Representative raw fluorescence traces showing glutamate-evoked calcium responses in control (**C**) and Spartin-DN-expressing (**D**) primary cortical neurons expressing jGCaMP7b at DIV14 in the presence of TTX. Quantification is shown in Fig. 3Q.

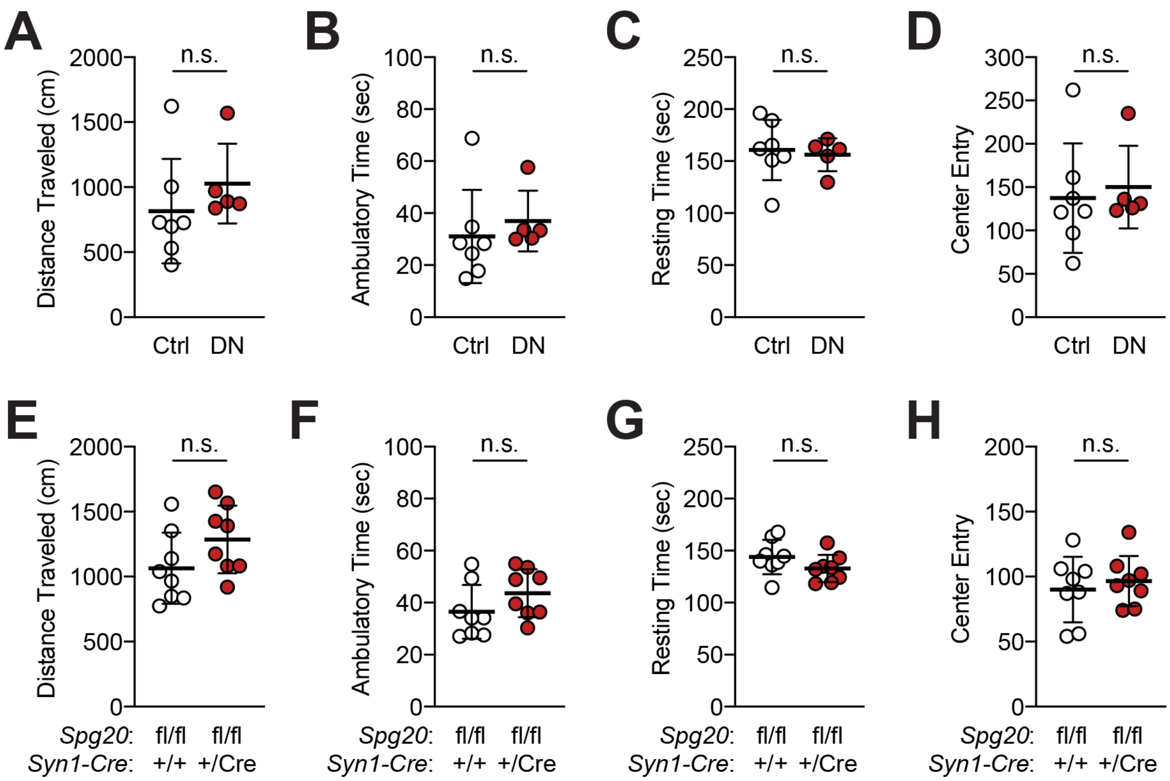


**Fig. S7. General locomotion of Spartin-DN-expressing mice and neuron-specific Spg20 conditional KO mice.**
(**A** to **D**) Open field data from neuron-specific Spartin-DN-expressing mice. No significant differences between control and Spartin-DN-expressing mice in distance traveled (**A**), ambulatory time (**B**), resting time (**C**), and center entry (**D**). Mean ± s.d.; $n$ = 7 control and 5 Spartin-DN mice. Statistical significance was determined by Mann–Whitney tests (n.s., non-significant). (**E** to **H**) Open field data from neuron-specific Spg20 conditional KO mice. No significant differences between control and Spg20 conditional KO mice in distance traveled (**E**), ambulatory time (**F**), resting time (**G**), and center entry (**H**). Mean ± s.d.; $n$ = 8 mice per genotype. Statistical significance was determined by Mann–Whitney tests (n.s., non-significant).
